# It’s not the spoon that bends: Internal states of the observer determine serial dependence

**DOI:** 10.1101/2023.10.19.563128

**Authors:** Ayberk Ozkirli, David Pascucci

## Abstract

Traditional views suggest that human perception handles uncertainty using optimal strategies. For instance, when prior stimuli are more reliable than current ones, perceptual decisions rely more on the past, leading to stronger serial dependence. Here, we report findings that challenge this view. We asked human observers to reproduce the average orientation of an ensemble of stimuli under varying stimulus uncertainty. Contrary to optimal strategies, we found that serial dependence is stronger when prior stimuli are more uncertain. We hypothesize that fluctuations in stimulus uncertainty may influence internal states of observers, such as participants expectations about uncertainty and beliefs about their own performance. A striking finding is that manipulating these internal states through rigged feedback can yield drastic effects on serial dependence, even when external input (i.e., stimulus uncertainty) remained constant. Our findings suggest that phenomena like serial dependence can be better understood by considering internal states of the observer, beyond fixed computations and optimal strategies.

## Introduction

No one likes uncertainty. Yet, uncertainty is not only a persistent aspect of our environment, but also an extremely volatile one, evolving swiftly and sometimes with dramatic consequences. Imagine a pilot navigating through rapidly changing weather conditions, facing the risk of making the wrong decisions by relying on suddenly unclear visual cues.

According to traditional models of perception and decision-making, the brain deals with uncertainty in a near-optimal and Bayesian way^1–9^. For example, when a stimulus is unclear or ambiguous, the brain leverages more reliable sources of information, such as prior experiences, to guide its interpretation^9–12^. A central tenet of these models is that uncertainty is computed from each stimulus, and then multiple stimuli is integrated, with a greater weight to the more reliable ones^2,4,10,13,14^. Models of this kind, often described within the framework of the ‘ideal observer’^15^, can explain performance in a variety of perceptual tasks, suggesting that, in most cases, humans can effectively combine information from multiple stimuli to reduce uncertainty in an optimal way.

However, not all human behavior strictly adheres to Bayesian and optimality principles^16–19^. Recent research on serial dependence, the phenomenon whereby perceptual decisions are biased toward previous stimuli^20,21^, provides a clear example. Ideal observer models predict that serial dependence is stronger when the previous stimulus is more reliable than the current one, as the optimal strategy is to rely more on more reliable stimuli^10^. Nonetheless, existing results are rather mixed. Some studies show that only the present uncertainty, not the past, influences serial dependence^22–24^, while others report increased serial dependence when the uncertainty on the previous trial is high, rather than low^25,26^. In sum, human perceptual decisions do not seem to weigh prior stimuli in the same way as ideal observers would.

The reasons for such discrepancies can be manifold, as multiple factors contribute to serial dependence. These include the interplay between attractive and repulsive forms of serial dependence^27–30^, modulations due to higher-level cognitive functions, such as confidence, attention, working-memory and expectations^20,27–29,29–34^, among others. These factors may all affect the way uncertainty is handled in perceptual decision-making. For instance, a change in stimulus uncertainty might be internally perceived as a change in the state of the world^32^ —e.g., an uncertain stimulus is likely to be followed by another uncertain one. Consequently, individuals may adjust how they integrate information over time, particularly when experiencing moments of high uncertainty. These changes in integration strategies may ultimately impact the strength of serial dependence, beyond the predictions offered by ideal observer models alone. The contribution of these factors, such as participants expectations about uncertainty and beliefs about their own performance —termed here as ‘internal states’— cannot be adequately captured by models predicting serial dependence by relying exclusively on stimulus uncertainty.

In this study, we focused on the role of internal states in serial dependence and how they may, at least partially, explain patterns in behavior that deviate systematically from ideal observer models. In a first step, we tested how transitions between different levels of uncertainty affect serial dependence in an orientation averaging task, where participants reproduced the average orientation of an ensemble of stimuli (Figure 1A). In two experiments, we found that serial dependence increased when the previous stimulus was uncertain (e.g., an ensemble with a wider distribution of orientations) and was the largest when both the current and previous stimuli were uncertain—a pattern nearly opposite from that predicted by the ideal observer. To explain this behavior, we hypothesized that the persistence of high uncertainty across consecutive moments induces an internal state that promotes serial dependence—a hypothesis formalized in an accompanying model. Building on this hypothesis, we predicted that, even under constant stimulus uncertainty, changes in internal states could still influence serial dependence. To test this prediction, we conducted a final experiment where we manipulated performance feedback. Strikingly, serial dependence was observed only when participants believed their performance was poor due to manipulated feedback, despite the level of stimulus uncertainty, performance, and the feedback received on the most recent trial remaining constant.

**Figure 1.**
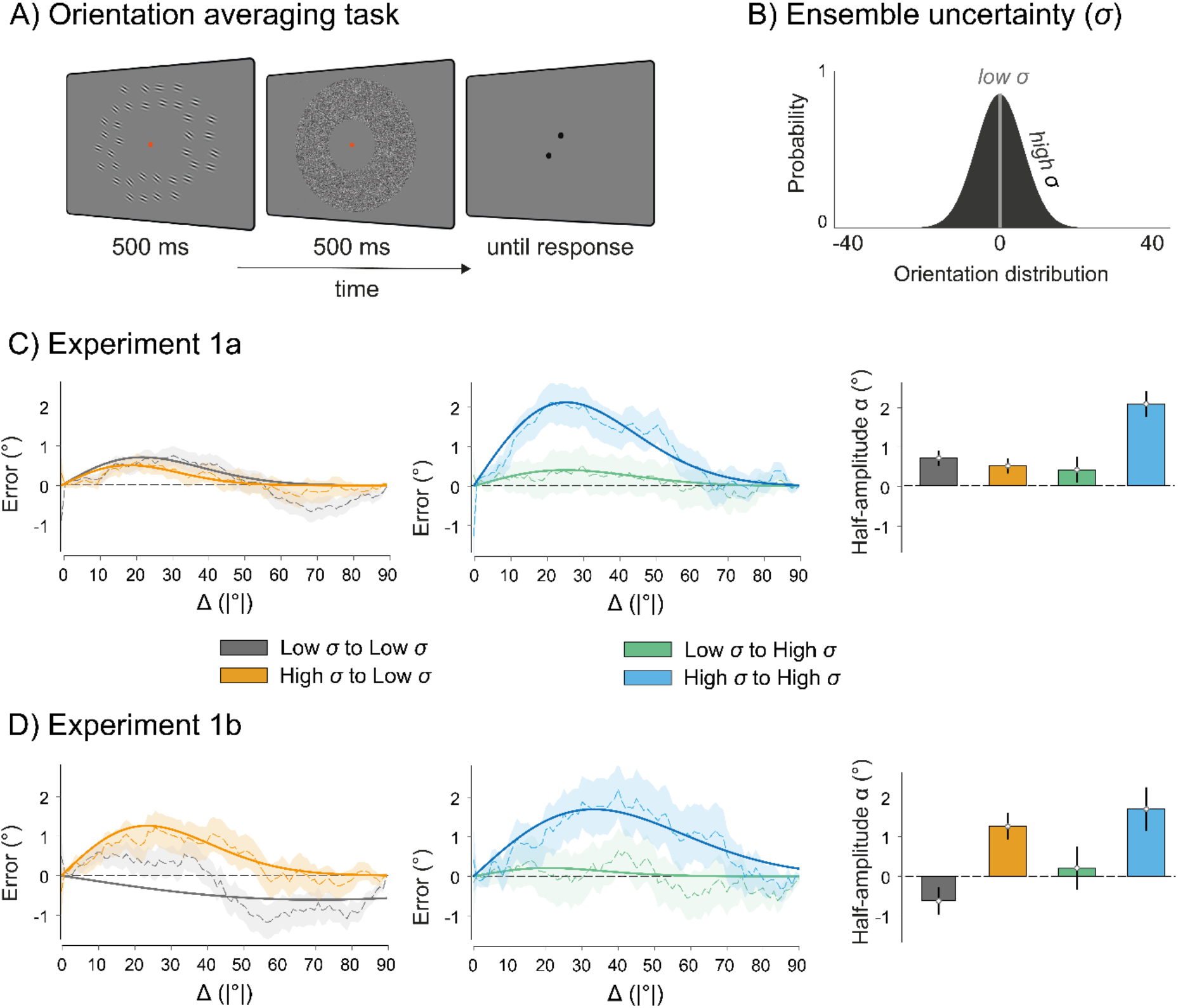
Main paradigm and results of Experiment 1a-b. A) Participants reproduced the average orientation of an ensemble of 36 Gabors shown in the periphery of the visual field (see Methods). B) We manipulated uncertainty by intermixing ensembles with high and low orientation variability (e.g., standard deviation [σ] of the orientation distribution, σ = 8° vs. 0° in Experiment 1a, σ = 15° vs. 5° in Experiment 1b). C) The first panel shows serial dependence in the high-to-low (in orange) and low-to-low conditions (in grey). The second panel shows serial dependence in the high-to-high (in blue) and low-to-high conditions (in green). A circular running average (sliding window size of 35°) is computed on the folded adjustment errors of all participants as a function of the absolute difference in average orientation between the previous and current ensemble (Δ; |previous minus current|, see Methods). Shaded areas are 1 standard deviation of the running average. The curves represent the average first Derivative of Gaussian (δoG) fit across participants. The bar plots in the third panel show the average half-amplitude parameter (α) of δoG fits across participants, for each combination of prior and current uncertainty. Error bars represent standard errors of the mean, corrected for within-subject repeated-measures designs^36^. D) Results from Experiment 1b where the two σ used were 15° and 5°. The color coding is the same as in C.

**Figure 2.**
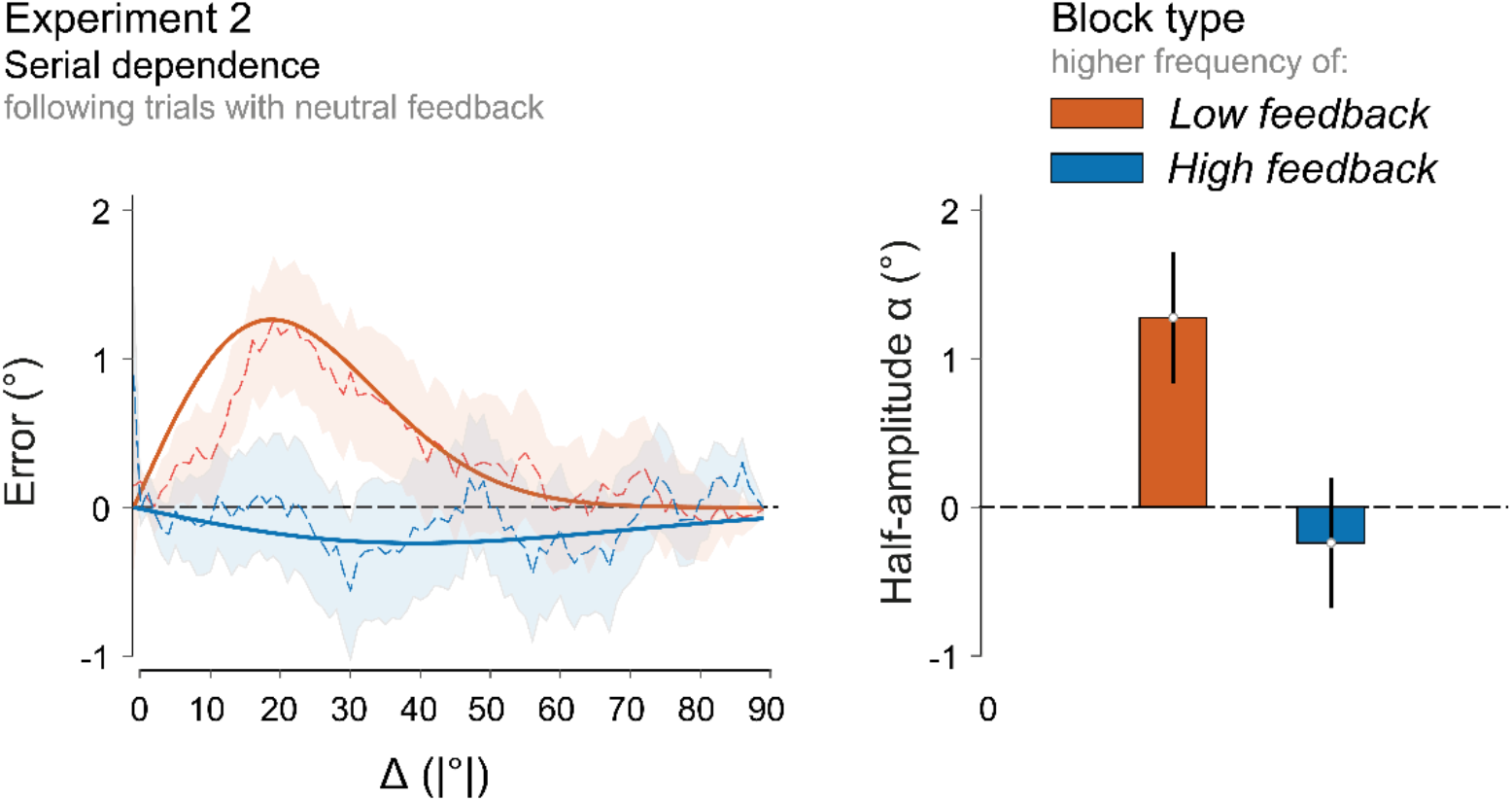
Results of Experiment 2. Serial dependence in the two types of blocks (red: *low feedback*, blue: *high feedback*). *Low feedback* blocks contained a higher number of poor performance feedback trials. *High feedback* blocks contained more good performance feedback trials. In both blocks, the two types of feedback were intermixed with equal frequency of neutral feedback trials (see Methods). Serial dependence was assessed on trials following neutral feedback to mitigate any potential impact of feedback on the immediately preceding trial. The graph on the left depicts moving averages of folded errors and the best fitting δoG, similar to Figure 1. Bar plots on the right represent the average of the δoG half-amplitude parameter (α) across participants in the two conditions. Error bars represent standard errors of the mean. Note that performance (i.e., the standard deviation of adjustment errors) remained constant across conditions, despite evidently different levels of serial dependence (see main text).

In summary, our findings suggest that phenomena like serial dependence are largely flexible, contingent upon internal states triggered by the need to mitigate uncertainty, and cannot be reduced to ideal observer models that rely solely on stimulus parameters.

## Results

### The effect of transitions in stimulus uncertainty on serial dependence

In serial dependence, perceptual decisions are biased towards the stimulus shown on the preceding trial^21,35^. According to an ideal observer, the bias should be larger when the current stimulus is more uncertain than the previous one, and reduced when the current stimulus is more reliable^10^ (see Figure S3A in Supplementary Material).

To test these predictions, we asked sixteen observers to reproduce the perceived average orientation of an ensemble of 36 Gabor stimuli (Figure 1A). On randomly interleaved trials, the ensemble orientations were either drawn from a normal distribution with a standard deviation (σ) of 8° (*high* σ) or all identical (*low* σ = 0°). These two conditions carried different degrees of uncertainty, as reflected in performance, with a larger standard deviation of adjustment errors in the *high* σ (12.56 ± 2.69°) compared to the *low* σ condition (6.77 ± 1.18°; *high* vs. *low*: t(15) = 12.31, *p* < .001, Cohen’s *d’* = 3.08, paired t-test).

We estimated the strength of serial dependence using the half-amplitude parameter from a δoG fit (α, see Methods), considering all possible combinations of current and past uncertainty, in a full-factorial design. A repeated-measures ANOVA with factors ‘Uncertainty on the preceding trial’ (*low* vs. *high*) and ‘Uncertainty on the current trial’ (*low* vs. *high*) revealed no effect of the current trial uncertainty (F(1, 15) = 2.22, *p* = .16, 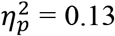), but a significant main effect of the uncertainty on the preceding trial (F(1, 15) = 5.21, *p* = .037, 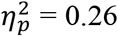) as well as a significant interaction (F(1, 15) = 21.19, *p* < .001, 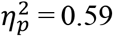). The interaction was driven by the serial dependence occurring when both the previous and current stimuli were of high uncertainty (the *high-to-high* condition), which was significantly larger when compared to all the other conditions (*high-to-high* vs. average of the other conditions: t(15) = 3.837, *p* = .0016, Cohen’s *d’* = 0.959; see Figure 1C).

This finding was inconsistent and even nearly opposite to what would be expected by the ideal observer with classic optimal integration (see Supplementary Material). In other words, observers did not exhibit stronger biases when the prior stimulus was more reliable; instead, they exhibited the strongest bias toward past stimuli when uncertainty was high in both the current and previous trials.

One aspect worth noting is that the interaction term was significant: when the current uncertainty was low, we did not find any effect of the previous trial uncertainty (Figure 1C, first panel). This may be attributed to ceiling performance and weak serial dependence in the low uncertainty condition, where all stimuli shared the same orientation. Another possibility is that *low* σ stimuli also carried stronger repulsive effects that contrasted attractive serial dependence^27–29,37,38^, since repulsive biases are often observed after well-visible stimuli that can be represented with high precision^39–41^.

To address these concerns, we performed a follow-up experiment with 16 new participants, increasing the standard deviations (σ) in both the *high* (σ = 15°) and *low* (σ = 5°) uncertainty conditions. As in the first experiment, observers were less precise in the *high* σ (standard deviation of adjustment errors: 16.54 ± 4.90°) compared to the *low* σ condition (11.05 ± 2.78°; *high* vs. *low*: t(15) = 7.37, p < .001, *d’* = 1.84, paired t-test).

Upon increasing σ in both conditions, a repeated-measures ANOVA revealed a significant effect of the preceding trial (F(1, 15) = 11.698, *p* = 0.0038, 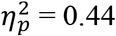), but no significant effect of the current trial uncertainty (F(1, 15) = 0.796, *p* = 0.386, 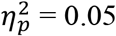) or the interaction (F(1, 15) = 0.217, *p* = .65, 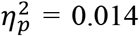). Therefore, when uncertainty was increased in both conditions compared to Experiment 1a, serial dependence was once again overall stronger when the preceding stimulus was more uncertain, regardless of the current trial uncertainty. Further analyses indicated that these effects strengthen with an increasing number of consecutive high-uncertainty trials in the past (see Figure S2), thereby excluding the possibility that the observed differences stemmed from other confounding factors associated with the representation of average orientations on the immediately preceding trial (e.g., heightened repulsive adaptation-like aftereffects following more precise orientation representations).

The patterns observed in Experiment 1a-b, despite starkly contradicting the predictions of ideal observer models of serial dependence (see Figure S3A), can be more accurately captured through a straightforward yet effective conceptual extension of these models. Indeed, a model that incorporates the impact of sequences of uncertain stimuli on the observer internal states can effectively adjust the trial-by-trial tendency to integrate information over time, outperforming the fit of a simple ideal observer in both Experiment 1a and 1b (see *State-dependent model of serial dependence* in Supplementary Material).

The key concept here is that serial dependence is not solely determined by a fixed computation of uncertainty based on the stimulus; rather, it depends on internal states of the observer, that evolve as a function of the history of stimulus uncertainty. This led us to formulate a key prediction: even when stimulus uncertainty remains constant, internal states of an observer can entirely dictate the strength of serial dependence. To directly test this prediction, we conducted a final experiment, which also served to definitively rule out any other explanation of the results based on differences between the physical parameters of the stimuli across conditions, such as adaptation-like effects and orientation bias.

### Internal states modulate serial dependence

We tested 24 new participants in a paradigm where we fixed the ensemble σ to 15° and introduced trial-by-trial feedback. The goal of the feedback was to manipulate internal states of the observer by changing their beliefs about their own performance, and thus the experienced uncertainty in different blocks of trials although the external uncertainty provided by the stimulus always remained constant (see Methods). To this end, we used two types of blocks: *low feedback* blocks, where observers received more frequent feedback indicating poor performance (“below average”), and *high feedback* blocks, where they received more frequent feedback indicating good performance (“above average”). In both types of blocks, low and high feedback trials were intermixed with an equal number of trials in which participants received neutral (“average”) feedback. Importantly, within each block, the feedback indicating poor or good performance still reflected the actual performance levels (see Methods). We focused on ‘block’ effects on serial dependence —i.e., modulations due to the overall frequency of *low* or *high feedback* in a block—and thus, to control for any effect of the specific feedback received on the most recent trial, we restricted our analysis to trials following neutral feedback, which was received with equal frequency in both block types.

Participants performed the task with comparable levels of performance between the two block types (standard deviation of adjustment errors: *low*: 13.73 ± 3.59°, *high*: 14.20 ± 3.44°, *low* minus *high*: t(23) = -1.65, *p* > .05, *d’* = -0.34). Strikingly, however, and in line with our predictions, serial dependence was significantly larger in the *low feedback* compared to the *high feedback* blocks (*low* minus *high*: t(23) = 2.44, *p* = .023, *d’* = 0.50). Thus, even though the stimulus uncertainty was fixed, and performance was comparable between conditions, manipulating internal states of the observer led to clear differences in serial dependence: under the internal belief of overall worse performance in a sequence of trials, the tendency to integrate current and prior stimuli increased.

## Discussion

In this study, we investigated the impact of uncertainty on serial dependence in perceptual decisions. Existing frameworks, based on ideal observer models, predict stronger serial dependence when prior stimuli are more reliable than current ones^10^. Contrary to this prediction, we observed an increase in serial dependence when prior stimuli were more uncertain. We hypothesized that this phenomenon could be due to internal states of the observer that modulate serial dependence beyond the mere effects of stimulus uncertainty. We directly tested this hypothesis in a final experiment, where we manipulated internal states, namely participants’ beliefs about their own performance in a block of trials, using feedback, and found drastic effects on serial dependence, even when the uncertainty in the stimulus remained constant.

Prior research has modeled serial dependence following the classic assumptions of optimal and Bayesian models^5,42,43^. These models posit that serial dependence is influenced by internal priors about external world stability and by the relative reliability between these priors and current sensory evidence^38,43,44^. However, empirical validation of these models has lacked a systematic assessment across all possible conditions of relative uncertainty. Through a full factorial design, we demonstrate that these models provide only a loose approximation, as they fail to account for a crucial aspect observed in two experiments (Experiments 1a-b): the increase in bias when the uncertainty of prior stimuli is high (see Figure 1).

We argue that our findings can be explained in the context of temporal expectations and adaptable internal beliefs^25,32^. Even in the absence of regularities and temporal structures, the brain constantly uses recent experiences to infer the current state of the environment^37,45–47^, a core notion of predictive theories of the brain^37,45,47,48^. These internal beliefs act as downstream, top-down predictions that are fed back and shape processing as early as in primary visual cortex^49^. Crucially, internal beliefs encompass not only specific stimulus features, but also the level of uncertainty associated with encountering a stimulus^32^. Thus, the occurrence of an uncertain stimulus may prompt a change in the internal states of the observer, leading to a greater tendency towards serial dependence as a means to mitigate uncertainty.

This interpretation is consistent with the notion that, while classic models assume a fixed structure of prior beliefs^38,43,44^, actual beliefs are dynamic and contingent on internal states that adapt and reset when changes in the environment are detected^25,50^. In contrast to static priors, indeed, we show that a simple extension of an ideal observer model, which incorporates internal states, provides a more accurate approximation of the data (see *State-dependent model of serial dependence* in Supplementary Material). This idea is reminiscent of Kalman filters that adapt and self-tune to changes in input data^42,51,52^, which may provide a promising modeling framework for future studies.

One may question the extent to which the findings of Experiment 1a-b could be accounted for by alternative explanations and more complex models. For instance, previous studies have highlighted the existence of both attractive and repulsive forms of serial dependence^27,28,37,38,53^. The interaction and relative prevalence of these opposing biases may be influenced by various factors, including the stimulus uncertainty, with lower uncertainty favoring repulsive effects. Although we cannot dismiss the potential role of a repulsive component, which might have countered attractive serial dependence in conditions of low uncertainty, our supplementary analyses revealed patterns that cannot be explained by this. Specifically, we observed that attractive serial dependence following an uncertain stimulus increased in strength as a function of the number of consecutive uncertain stimuli occurring before (see Supplementary Material for details). Moreover, Experiment 2, where stimulus uncertainty remained constant, provided clear and unequivocal evidence that internal states, independent of stimulus parameters, can significantly influence the strength and even the manifestation of serial dependence. This finding aligns with existing research that explores factors beyond the stimulus in understanding serial dependence, including confidence^54^, attention^20^, memory load^55^, time-on-task^56^, task demand^28,29,31,37,57^, stimulus predictability^58^, as well as internal (e.g., neural) sources of uncertainty^43^.

As noted earlier, other studies have also reported patterns that diverge from those predicted by simple ideal observer models. For example, Ceylan et al.^22^ found similar trends to our study, with the strongest serial dependence occurring in high-to-high uncertainty conditions. However, they did not observe significant differences between this and all other conditions, as we did. The reasons for this inconsistency are unclear and may stem from various factors, including differences in stimuli (single stimuli vs. ensembles) and the manipulation of uncertainty (e.g., changes in low-level features such as spatial frequency or noise vs. variability in ensembles), as well as differences in sensitivity (e.g., in^22^, the four combinations were tested only in one out of three blocks of trials, with a reduced number of trials per condition compared to the current study). Nevertheless, the presence of similar trends across multiple studies^22–26^ suggests that serial dependence cannot be solely explained by ideal observer and Bayesian models in their standard form. Building on this notion, recent research has used the term ‘partial Bayesian inference’ to describe the computations underlying serial dependence^24^. The term ‘partial’ implies the involvement of other contributing factors, and as demonstrated here, one such factor relates to the internal states of the observer, whose influence can nearly reverse the patterns predicted by ideal models.

Systematic deviations from ideal observer models, like the one reported here, do not necessarily imply that such models are incorrect, but highlight the need to move beyond simple and classic models that rely solely on basic aspects of the stimulus^32,33^. In the present context, differences in serial dependence can be found even without manipulating the stimulus parameters: what matters is the internal state of the observer. Similarly, in other domains like spatial vision, classic models that rely solely on local stimulus properties and feedforward processing may fail to explain phenomena like crowding and ensemble perception^59–62^.

In summary, we demonstrate that serial dependence is modulated by internal states of the observer, providing, at least a partial explanation of why the observed patterns typically deviate from the predictions of existing models. These results corroborate the view of serial dependence as a two-stage process, where effects are shaped not only by the stimulus but also by higher-level cognitive processes, in a recurrent hierarchical system where past experience plays a role at multiple interacting stages^28^.We argue that taking into account these aspects may not only lead to more accurate and generalizable predictions of serial dependence, but also to a better understanding of its functional roles, as flexible states of serial dependence may be more useful than a hard-wired and fixed mechanism. Integrating prior and current stimuli can be advantageous, particularly when anticipating moments of uncertainty in perception. However, integration may not be compulsory, especially when we can rely on our ability to discriminate between changes^58^.

## Methods

### Ethics statement

The study was approved by Commission cantonale d’éthique de la recherche sur l’être humain (protocol number 2021-02270) under the Declaration of Helsinki^63^, except for preregistration.

### Apparatus

Stimuli were presented on a gamma-corrected VG248QE monitor (resolution: 1920 x 1080 pixels, refresh rate: 120 Hz) and were generated with custom-made scripts written in MATLAB (R2013a) and the Psychophysics Toolbox^64^, on a Windows-based machine. Participants sat at approximately 57 cm from the computer screen, with the head held stable on a chin rest. The experiments were performed in a dark room.

### Participants

62 healthy participants (20 in Experiment 1a, 17 in Experiment 1b; age range of 18-37 years; 5 females in Experiment 1a, 9 females in Experiment 1b, and 15 females in Experiment 2), from the EPFL and the University of Lausanne, took part in the study for a monetary reward (25 CHF/hour). Participants were naïve to the purpose of the experiments. Written informed consent was collected from all participants beforehand.

### Stimuli and procedures

The stimuli and procedure are depicted in Figure 1. Each trial started with a dark red fixation spot appearing for 300-1000 ms (randomly sampled in steps of 100 ms). Participants were then presented with a display of 36 oriented Gabor stimuli for 500 ms (peak contrast: 25% Michelson, spatial frequency: 2 cycles/°, Gaussian envelope: 0.225°, phase = 0.25). Gabor stimuli were presented on a gray background (62.66 cd/m^2^) and positioned inside an annulus centered at the fovea (internal radius of 4.825°; external radius of 9.175°). No stimuli were presented in the central part of the display to avoid systematic biases toward the center during mean orientation estimates^62,65^. Inside the annulus, Gabor stimuli were arranged at equidistant locations (approximately 2.2° with a 1° random jittering) inside a 12x12° grid. To prevent after-effects, a noise mask was presented within the same annulus for 500 ms immediately after the Gabor ensemble. Following a blank interval of 1000 ms, participants reproduced the perceived average orientation by adjusting a response tool. The response tool consisted of two symmetric dots, each with a diameter of 0.25°, positioned approximately 1.5° apart, marking the endpoints of an imaginary line. The orientation of the response tool was initialized randomly, and participants were instructed to rotate the imaginary line using the mouse and confirm their response with a left click.

On each trial, the orientation of the Gabor stimuli was drawn from a normal distribution with the mean randomly selected over the full 0-179° orientation range, in steps of 1°. Experiment 1a involved two randomly intermixed conditions in which the standard deviation (σ) of the ensemble’s orientations was either 0° (*low* σ) or 8° (*high* σ). In Experiment 1b, σ was increased to 5° (*low* σ) and 15° (*high* σ). In Experiment 2, it was fixed at 15°.

In Experiment 2, each trial was followed by written feedback, shown at the center after 1 s from the response: “below average”, “average”, or “above average”. To establish the criteria for assigning feedback, we first determined an error threshold based on the mean absolute error (11°) estimated from the group of participants in Experiment 1b, in the condition with σ = 15°. The frequency was manipulated across different blocks. In the *low feedback* blocks, we assigned “below average” feedback on 75% of the trials where the error exceeded the threshold, and “average” feedback on the remaining 25% of those trials. Conversely, for trials in which the error was below the threshold, we assigned “above average” feedback on only 25% and “average” feedback to 75% of them. The opposite assignment pattern was applied in the *high feedback* blocks. In other words, “below average” feedback was given on only 25% of trials where the error was above the threshold, while “average” feedback was assigned to the remaining 75% of those trials. For trials with errors below the threshold, “above average” feedback was assigned to 75% and “average” feedback to 25% of them. As a result of this manipulation, participants were more likely to receive “below average” (indicating poorer performance) in the *low feedback* blocks and “above average” (indicating better performance) in the *high feedback* blocks, despite their actual performance potentially remaining consistent.

An important aspect of this manipulation was that the frequency of trials with “average” feedback was the same in both blocks. This allowed us to examine serial dependence based on the overall block condition (*low* vs. *high feedback*) focusing exclusively on trials following neutral feedback, thus controlling for the specific feedback received on the preceding trial.

To effectively manipulate internal states, it was also important that the feedback was perceived as reliable and reflective of the actual performance. According to our debriefing with participants at the end of Experiment 2, 23 out of 25 of them reported having had poorer performance in some blocks, and 22 of those reported that this was due to difficulty difference across blocks (even though the stimulus σ was constant).

All participants received verbal and written instructions at the beginning of each experiment and completed a series of practice trials under the experimenter’s guidance. The data from the practice trials were not included in the analysis. In total, participants completed 400 trials in all three experiments. The trials were divided into 10 blocks for Experiment 1a and 1b, and 4 for Experiment 2. The *high* and *low feedback* conditions alternated across the blocks, with the order of the conditions counterbalanced among participants. Participants were instructed to keep their gaze fixed at the center of the screen throughout each block. All experiments lasted about 1 hour.

### Preprocessing of adjustment errors

Adjustment errors were computed as the acute angle between the response and the stimulus on each trial. We first excluded trials where errors were larger than 1.5 interquartile ranges above the upper quartile or below the lower quartile (separately for each experimental condition) and reaction times larger than 10 seconds. The first trial of each block, as well as trials following a removed trial, were also excluded. Overall, less than 10% of trials were removed in all experiments. In both Experiment 1a and 2, one participant was excluded due to a large overall negative bias (< -5°). In addition, participants were excluded if their biases in a specific condition of interest were 1.5 interquartile ranges larger than the upper quartile or smaller than the lower quartile within that condition (3 in Experiment 1a and 1 in Experiment 1b). The final analyses included 16 participants in Experiment 1a, an independent group of 16 participants in Experiment 1b, and 24 participants in Experiment 2.

In addition to the data cleaning procedure mentioned above, we also evaluated the presence of orientation biases, another typical step in preprocessing orientation adjustment errors for serial dependence analysis^28^. Orientation biases—i.e., the non-uniform distribution of errors over the orientation space— can introduce noise and artifacts into the estimation of serial dependence. Given that orientation biases may vary depending on uncertainty^66^, we examined their presence and the necessity for their removal in each condition of Experiments 1a-b, where uncertainty varied (see Supplementary Material). We found a clear orientation bias only in the condition with σ = 0° (Experiment 1a, see Figure S1). This finding aligns with previous research indicating that while orientation bias is typically apparent with a single stimulus (or all identical stimuli), its magnitude can vary significantly and be substantially reduced in the context of more heterogeneous ensembles^66^. The bias was removed by taking the residuals of a *malowess* function in MATLAB. Note that even without this additional step, the results remained consistent (see Supplementary Material).

### Analysis of serial dependence

In Experiment 1a and 1b, we separated the conditions based on the preceding and current stimulus uncertainty, which resulted in four categories: *low-to-low, high-to-low, low-to-high*, and *high-to-high*. We fitted the first derivative of a Gaussian function (δoG) to the errors as a function of the difference in orientation between the previous and current average orientation of the ensemble (Δ). The δoG has the following form:

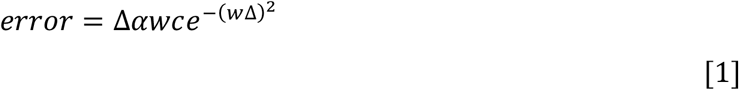

where *c* is a constant 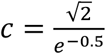 and *w* is the inverse of the curve width. The half-amplitude parameter α quantifies the deviation of the errors, in degrees, from the actual orientation, as a function of Δ: positive values of α indicate a systematic deviation of errors towards the orientation of the preceding stimulus, and negative values indicate a deviation away —i.e., repulsion. The parameter *w* was estimated using an aggregate participant for each condition^31,67^. Then, α was estimated for each participant and condition separately, using the respective aggregate *w* for that condition. For statistical purposes, repeated-measures ANOVA and t-tests were performed on the estimated α, depending on the experimental design and comparison of interest. For ANOVA, the effect sizes were reported in terms of partial-eta squares (0.01 to 0.059 indicating a small effect size, 0.06 to 0.139 indicating a medium effect size, and 0.14 or higher indicating a large effect size). For t-tests, we reported Cohen’s d (0.2 to 0.49 indicating a small effect size, 0.50 to 0.79 indicating a medium effect size, and 0.8 or higher indicating a large effect size).

## Supporting information

Supplementary analyses

## Acknowledgments

This research was supported by funding from the Swiss National Science Foundation (grant no. PZ00P1_179988 and PZ00P1_179988/2 to DP). The funders had no role in study design, data collection, and analysis. The authors would like to thank Bianca Ribeiro Riccioppo and Zehra Merchant for helping with the collection of the data.

## Data availability

The data will be made available in online repositories upon publication.

## Author contributions

The authors contributed equally to this work.

