## Supplementary analyses for "It’s not the spoon that bends: Internal states of the observer determine serial dependence"

### Supplementary Material

#### Orientation bias removal

Orientation adjustment tasks are not only influenced by serial dependence but also by systematic biases due to the non-uniform representation of orientations. In particular, the 'orientation bias' is often observed, wherein responses are biased away from the cardinal orientations and towards the obliques[1]. Several studies have followed the approach of removing this bias from responses before analyzing serial dependence to avoid spurious effects or confounding factors. For instance, differences in orientation bias across conditions may artificially create differences in serial dependence[2].

To account for potential confounds related to orientation bias, we adopted a similar approach. We first assessed the presence of orientation bias in each condition of interest, recognizing that biases typically observed with single stimuli may differ substantially or be absent when ensembles with heterogeneous orientations are presented[3]. Indeed, we found a clear orientation bias only in the *low* σ (0°) condition of Experiment 1a, where all stimuli shared the same orientation. In contrast, biases were not evident in other conditions, and responses were centered around the actual average orientation (Figure S1). We therefore removed the orientation bias from the *low* σ condition of Experiment 1a only, by taking the residuals of a moving average LOWESS function (*malowess.m* in Matlab), with errors as the dependent variable and actual orientation (Theta) as the independent variable.

*Results*

The results reported in the main text are after orientation bias removal, however, the results were consistent even without the removal. That is, a repeated-measures ANOVA in Experiment 1a with factors 'Uncertainty on the preceding trial' (low vs. high) and 'Uncertainty on the current trial' (low vs. high) revealed no effect of the current trial uncertainty (F(1, 15) = 1.54, $p$ = .23, $\eta_p^2$ = 0.09), but a significant main effect of the uncertainty on the preceding trial (F(1, 15) = 6.86, $p$ = .019, $\eta_p^2$ = 0.31) as well as a significant interaction (F(1, 15) = 14.32, $p$ = .002, $\eta_p^2$ = 0.49). The interaction was driven by the serial dependence occurring when both the previous and current stimuli were of high uncertainty (the high-to-high condition), which was significantly larger when compared to all the other conditions (high-to-high vs. average of the other conditions: t(15) = 4.566, $p$ < .001, Cohen's $d'$ = 1.141).

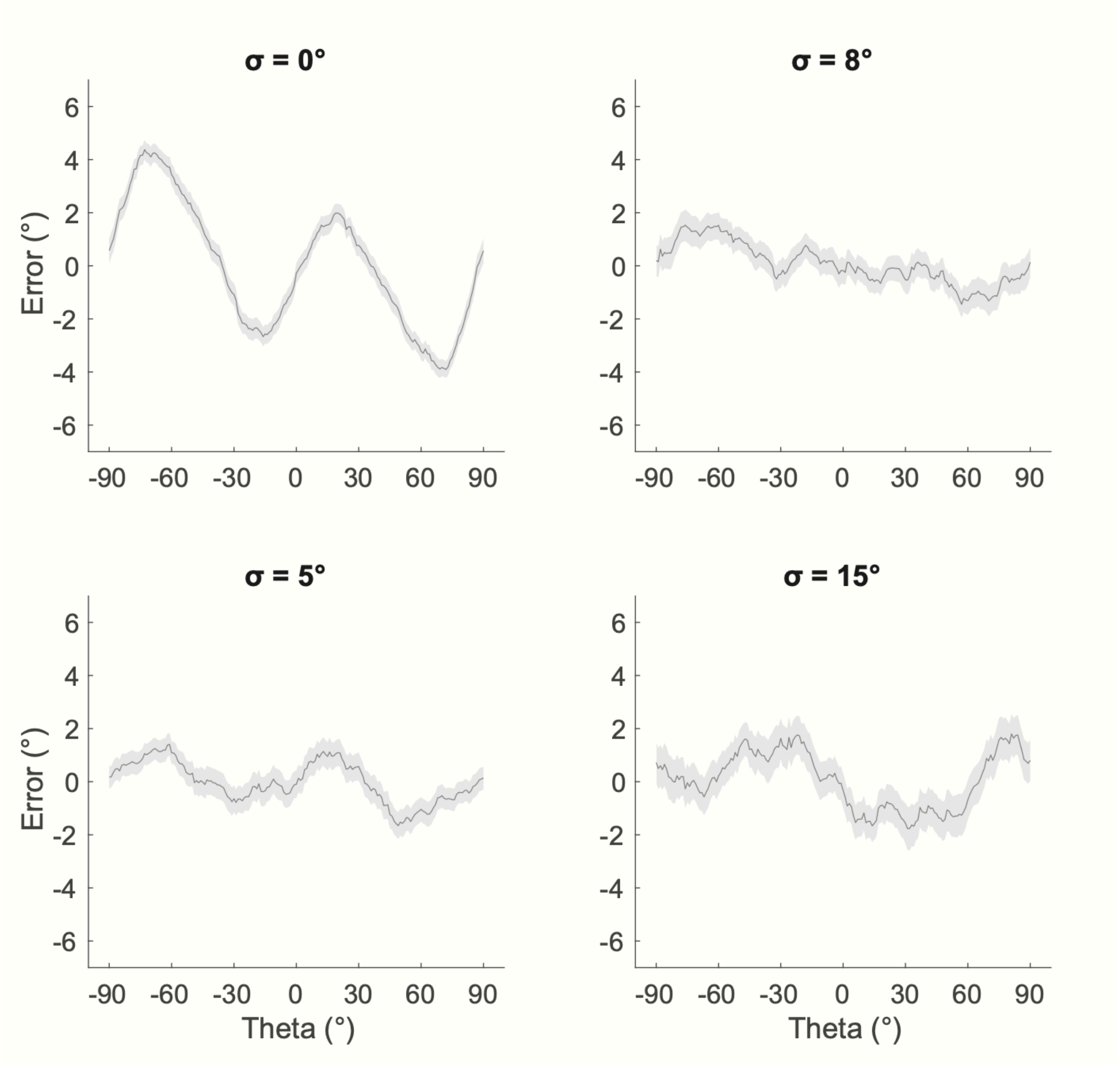


**Figure S1**. Orientation bias in the two uncertainty conditions of Experiment 1a (first row) and 1b (second row). The moving average of errors (y axis) is plotted as a function of the orientation shown in the present trial (Theta, y axis, 0° = vertical). The orientation bias is evident only in Experiment 1a, with σ = 0°, as the typical repulsion away from cardinals and towards obliques —i.e., orientations that deviate about 15° from the nearest cardinal (e.g., ±75° or ±15° in the x axis) are reported with larger errors, as shifted further away from cardinals. None of the other conditions show a similar pattern.

#### Sequence analysis

The results of Experiments 1a-b may also be attributed to the combined influence of additive repulsive and attractive effects exerted by prior stimuli, with varying degrees of strength across conditions[1]. For example, the weaker serial dependence in *low-to-high* trials could have been the result of stronger repulsive components induced by the low uncertainty in the preceding trial, as repulsion may tend to be more pronounced after precise and less ambiguous stimuli.

If the patterns observed were instead due to changes in internal states, triggered by the history of uncertainty, this should be evident in effects that extend beyond the immediately preceding trial. For instance, the effect of the previous stimulus uncertainty should also scale as a function of the uncertainty experienced several consecutive trials in the past. Such effect of more remote trials would rule out any confound due to the specific parameters of the stimulus and the related strength of repulsive components on the most recent trial.

To test this, we performed an exploratory analysis combining the data from both Experiment 1a and 1b (z-scoring the errors from each participant). We then separately analyzed conditions in which the previous trial was *high* or *low* σ. We created a 'sequence' variable indicating how many times in the past the same σ condition repeated (e.g., three consecutive trials with the same σ would be coded as '3'). We then fitted the errors in a restricted |Δ| range between 0° and 45°, where the relation with |Δ| approximates a linear trend, to a linear model of the form: *error* ~ |Δ|*σ**sequence* —i.e., errors were modeled via a 3-way interaction model considering the orientation difference with the previous trial (|Δ|), the σ on the previous trial, and how many times the same σ repeated before, as the main variables.

*Results*

The linear model revealed a main effect of |Δ| (coefficient = 0.0036±0.0012, $p$ = .0028), as well as a significant interaction between σ and |Δ| (coefficient = -0.0043±0.0017, $p$ =.01347) and a three-way interaction |Δ|*σ**sequence* (coefficient = 0.0021±0.0007, $p$ = .0031). Following the omnibus interaction, we evaluated the interaction between *sequence* and |Δ| separately for each σ condition, by splitting the model in two. The results revealed that the interaction |Δ|*σ**sequence* was mostly due to the fact that the positive relationship between errors and |Δ| —i.e., serial dependence— after trials with *high* σ, increased in strength as a function of the number of consecutive *high* σ trials before |Δ|**sequence* (coefficient = 0.0014±0.0005, $p$ = .0077, only considering *high* σ trials), while no effect of *sequence* was evident after *low* σ trials (only a significant effect of |Δ|, coefficient = 0.0036±0.0012, $p$ = .0025).

To depict this result, in Figure S2, we calculate a measure of the bias (average of folded errors within Δ range of 0-45°) for different values of *sequence* (1, 2 and >2). As evident, the more the repetitions of *high* uncertainty trials, the stronger the serial dependence effect, in line with our interpretation and the 'state-dependent' idea (see also State-dependent model of serial dependence below).

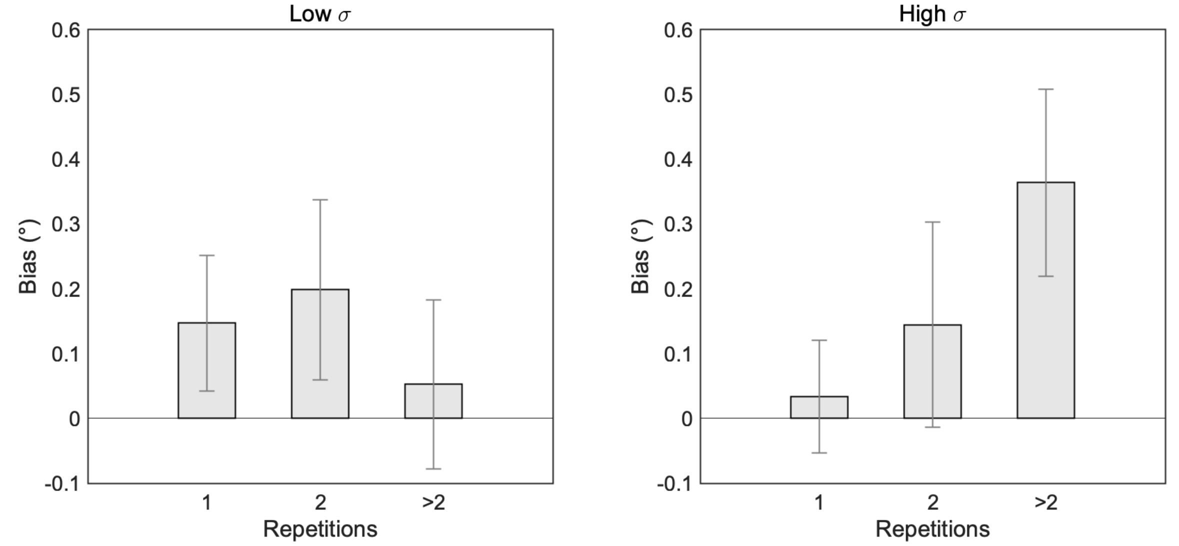


**Figure S2**. Serial dependence bias (positive means attraction towards the previous stimulus orientation) computed as a function of the σ on the preceding trial (*low* on the left plot, *high* on the right plot), and of the number of consecutive trials with the same σ condition presented before ('Repetitions', x axis). Serial dependence increased in strength with the number of trials in which the *high* σ condition repeated.

#### State-dependent model of serial dependence

Our interpretation of the results of Experiment 1a-b is that the uncertainty in a series of stimuli can lead to changes in internal states of the observer, such as expectations about consecutive moments of high or low uncertainty, which eventually interact with the strength of serial dependence (see Figure S3B). For example, the occurrence of an uncertain stimulus may increase the tendency to integrate prior stimuli into current perceptual decisions due to the expectation of a transition into a more uncertain state of the environment.

As a proof of concept, we here formalized this idea into an extended version of the ideal observer model, which we called the 'state-dependent model'. We then compared its ability to approximate the data of Experiment 1a-b against the classic ideal observer (see Figure S3).

In the ideal observer model, serial dependence results from how prior and current stimuli are relatively weighted[4–8]. Weights depend on the relative uncertainty of stimuli and on prior expectations. For example, priors may be composed of a mixture of a Gaussian-like probability distribution centered on the previous stimulus orientation, and a uniform distribution covering the whole orientation space[8]. The ratio between these two distributions determines the overall tendency to weight prior stimuli —i.e., when the uniform is the dominant component, the prior on past stimuli is flat, in a Bayesian sense, and there is no effect of prior stimuli.

We implemented the ideal observer model using a population coding approach[1]. In this approach, the representation of the average orientation is approximated via a probabilistic distribution over the orientation space. The distribution peak reflects the mean orientation of the ensemble, and the width is related to the uncertainty, parametrized by the mean and concentration parameter of a Von Mises distribution:

$$f_i(\theta) = e^{(k(\cos(\theta-\varphi_i)-1))}, \qquad [2]$$

where $f_i(\theta)$ is the response associated with the $i$-th orientation $\theta$ in the 0-179° orientation space, $k$ is the concentration parameter related to the width and $\varphi_i$ is the preferred orientation of each coding unit of the population. By normalizing $f_i(\theta)$ by its sum, the population response is represented as a probability distribution. In a Bayesian framework, eq. [2] and the resulting distribution describes the stimulus likelihood (e.g., $p(m|\theta)$, with $m$ = measurement)[5].

In a straightforward implementation of the ideal observer, the posterior $p(\theta|m)$ on each trail $t$ is the result of the combination between the likelihood and the prior —i.e., a representation of previous events (posterior) propagating from the trial $t-1$:

$$p(\theta^t|m^t) \propto p(m^t|\theta^t) * p(\theta^{t-1}|m^{t-1}). \qquad [3]$$

In serial dependence, changes in the uncertainty associated with each stimulus can be incorporated by varying $k$ in eq. [2] as a function of the ensemble standard deviation. For ease of interpretation, we define $\sigma_{low}$ and $\sigma_{high}$ as the two standard deviations of a wrapped normal distribution corresponding to two Von Mises concentration parameters $k$, and reflecting the ensemble's σ under the two levels of uncertainty. In line with previous work[8], $p_{past}$ is modeled as a weighted combination of the posterior distribution on trial $t-1$ and a uniform distribution:

$$p_{past}(\theta^{t-1}|m^{t-1}) = w * \left(p(\theta^{t-1}|m^{t-1})\right) + (1-w) * U(0{,}179), \qquad [4]$$

where $U(0{,}179)$ is a uniform prior with equal probability at each orientation and $w$ is a parameter determining the relative weight of the two components. It is straightforward to note that $w$ regulates the strength of the effect of prior stimuli: when $w = 0$ the prior is uniform, and history has no effect.

When $w > 0$, and $\sigma_{low}$ and $\sigma_{high}$ are incorporated, eq. [3-4] shifts the mean of the posterior towards the prior, by an amount that depends on the relative σ of the prior and the likelihood, as in classic optimal integration schemes:[4,8] the smaller the σ on the preceding trial, the larger the shift. Using the circular mean of the posterior as the final estimate of the orientation on each trial, one can observe systematic biases, compatible with the predictions of an ideal observer model of serial dependence (Figure S1).

In the model described above, two factors modulate serial dependence: 1) the ratio between the prior centered on past stimuli and the uniform one; 2) the relative uncertainty (width) of past and current stimulus representations. Standard models assume that the former, the ratio, is fixed and therefore serial dependence is only determined by the relative uncertainty between stimuli, which is computed independently for each stimulus. By relaxing this assumption, we let the parameter $w$ vary as a function of the σ on the preceding trial —i.e., the ratio of the prior about previous stimuli over the uniform prior changed depending on the stimulus history and the associated uncertainty. In the state-dependent model, $w$ was considered a free parameter, functioning as an internal-state-dependent weight that was updated on each trial based on the value of σ.

We compared the ideal observer and the dynamic observer models by feeding them with the single-trial data of Experiment 1a and 1b. Note that the ideal observer has three free parameters ($\sigma_{low}$, $\sigma_{high}$, $w$) whereas the dynamic observer model has four ($\sigma_{low}$, $\sigma_{high}$, $w_{low}$, $w_{high}$). To account for this difference, we compared the two models using the Akaike Information Criterion (AIC)[9] which penalizes the number of parameters and is estimated using the following formula:

$$AIC = n_{trials} * log\left(\frac{\sum\left(errors_{predicted} - errors_{observed}\right)^2}{n_{trials}}\right) + 2 * n_{parameters}$$

[5]

A difference in AIC between two models (ΔAIC) larger than 2 is typically considered strong evidence in favor of the model with the lower AIC[10].

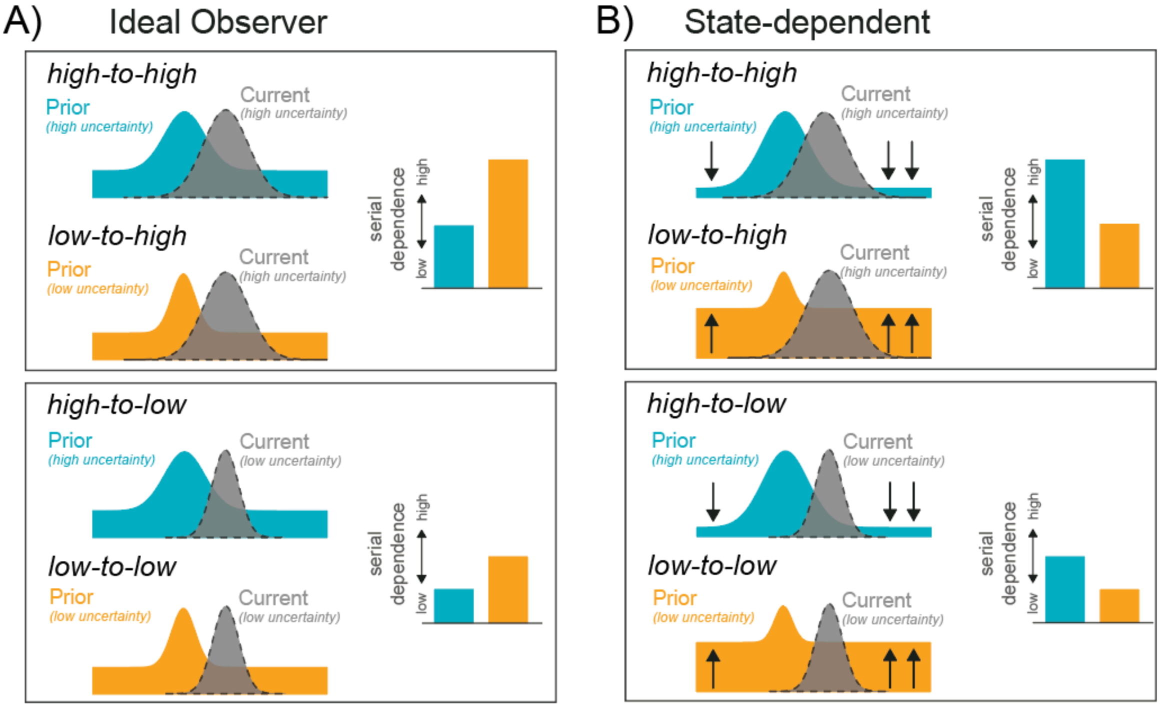


**Figure S3**. A-B) Two models of the effects of uncertainty in serial dependence. In the ideal observer model (A), prior and current stimuli are integrated depending on their relative uncertainty, with the more reliable stimulus always receiving larger weights. In such a model, serial dependence is always stronger when the previous stimulus is more reliable than the current one. The prior is here composed of a mixture of a uniform distribution and a normal distribution centered on the previous stimulus (blue and orange distributions, see Methods below for further details). Note that the strength of the prior here is only determined by the uncertainty in the stimulus, whereas the uniform component remains fixed. In the 'state-dependent' model (B), stimuli are integrated depending on their relative uncertainty as in the ideal observer, but the overall tendency towards integration (here modeled as the ratio between the uniform and normal prior) varies as a function of internal states: when an uncertain stimulus is encountered, the weight of the uniform component decreases and the tendency towards serial dependence increases. These changes are highlighted by the black arrows. The four boxes in A-B) depict all four combinations of prior and current uncertainty tested (see main text).

#### *Results*

The state-dependent model provides a more accurate representation of the actual patterns of serial dependence (see the bars in Figure S3 for the qualitative predictions of each model), outperforming the classic ideal observer model (difference in Akaike Information Criterion between the state-dependent and ideal observer models [$\Delta AIC_{Exp1a}$] = 8.28 (9.52 after orientation bias removal in Exp1a *low* uncertainty condition), and [$\Delta AIC_{Exp1b}$] = 4.26, where $\Delta AIC > 2$ favors the state-dependent model).
